# Astrocyte-to-microglia communication via Sema4B-Plexin-B2 modulates injury-induced reactivity of microglia

**DOI:** 10.1101/2023.11.07.565986

**Authors:** Natania Casden, Vitali Belzer, Abdellatif Elkhayari, Rachid Elfatimy, Oded Behar

## Abstract

After central nervous system injury, a rapid cellular and molecular response is induced. This response can be both beneficial and detrimental to neuronal survival in the first few days and increases the risk for neurodegeneration if persistent. Semaphorin4B (Sema4B), a transmembrane protein primarily expressed by cortical astrocytes, has been shown to play a role in neuronal cell death following injury. Our study shows that after cortical stab wound injury, cytokine expression is attenuated in Sema4B knockout mice and microglia/macrophage reactivity is altered. *In vitro*, Sema4B enhances the reactivity of microglia following injury, suggesting astrocytic Sema4B functions as a ligand. Moreover, injury-induced microglia reactivity is attenuated in the presence of Sema4B knockout astrocytes compared to heterozygous astrocytes. *In vitro*, experiments indicate Plexin-B2 is the Sema4B receptor on microglia. Consistent with this, in microglia/macrophage-specific Plexin-B2 knockout mice, similar to Sema4B knockout mice, microglial/macrophage reactivity and neuronal cell death are attenuated after cortical injury. Finally, in Sema4B/Plexin-B2 double heterozygous mice, microglial/macrophage reactivity is also reduced after injury, thus supporting the idea that both Sema4B and Plexin-B2 are part of the same signaling pathway. Taken together, we propose a model in which following injury, astrocytic Sema4B enhances the response of microglia/macrophages via Plexin-B2, leading to increased reactivity.

**Significance statement:** In this study, we show that in the brain cortex, Sema4B, a protein mainly expressed by astrocytes, plays a crucial role in enhancing the reactivity of microglia/macrophages via Plexin-B2. These findings reveal new molecular signaling instigated by astrocytes toward microglia/macrophages in the context of central nervous system (CNS) injury, shedding new light on the complex interplay between astrocytes and microglia/macrophages. Taken together, our findings suggest that targeting the Sema4B/Plexin-B2 pathway could be a promising therapeutic approach for reducing microglia reactivity and improving the adaptive response in the context of CNS injury.

## Introduction

Brain injury results in immediate damage and death to cells at the point of impact. As a result, there is an extracellular release of a variety of ions, molecules, and proteins termed damage-associated molecular patterns (DAMPs) (1). These molecules induce neuroinflammation, a response aimed at restoring tissue homeostasis. Although the inflammatory response is critical to limit damage, it has also been reported to exacerbate damage, including neuronal loss (2). Following this acute phase, neuroinflammation may persist and increase the probability of developing a neurodegenerative disease (3). Two of the major types of cells immediately activated by DAMPs are microglia and astrocytes.

Microglia rapidly responds to injury with dramatic morphological transformation (1). Nevertheless, microglial activation is not uniform in nature and depends in part on the distance from the injury (4). In their activated state, microglia can serve diverse beneficial functions essential to neuronal survival (5). However, activated microglia can also induce detrimental neurotoxic effects (6).

Astrocytes also respond to injury by a process called reactive astrogliosis which is characterized by changes in gene expression, morphology, and proliferation (7). Ablation of reactive astrocytes exacerbates cortical and spinal cord injuries (8). However, manipulating different signaling pathways in reactive astrocytes has shown that astrogliosis can increase or decrease neuronal survival depending on the specific pathways. For example, deletion of STAT3 in astrocytes results in increased neuronal death and more limited recovery following CNS injury (9), while inhibition of NF-κB, CCL2, or CXCL10 in astrocytes improves recovery (8). It thus appears that activation of astrocytes and microglia after injury induces opposing signaling pathways that together determine the degree of neuroinflammation and neuronal cell death. Thus, finding new molecular modulators of the brain response to injury may have the potential to affect neuroinflammation and neuronal cell death. One such possible candidate is Sema4B.

Sema4B is a member of the type 4 semaphorins, an evolutionarily conserved family that commonly acts as ligands that bind directly to plexins ((10). We have previously shown that Sema4B is expressed by cortical astrocytes and in its absence, the astrocyte activation profile is altered, and proliferation is reduced following cortical injury (11). We also showed that neuronal cell death 24h and 72h after a stab wound injury is reduced in these Sema4B knockout mice (12). These results imply that Sema4B affects the central nervous system’s response to injury. However, the mechanism by which Sema4B affects this response is not yet clear.

## Results

### Expression of Sema4B in the mouse brain

To test which cell populations express Sema4B in the cortex, we used double *in-situ* hybridization with probes for Sema4B, CX3CR1 (microglia/macrophage marker), and Sox9 (astrocyte marker). Most cells in the cortex expressing Sema4B are also Sox9 positive while only a small minority of Sema4B positive cells are also positive for Cx3Cr1 (Figure 1a, b, c). Additionally, most astrocytes express Sema4B (85.69% +/− 5.59) while only a minority of the microglia/macrophages express Sema4B (10.66% +/− 2.1).

**Figure 1:**
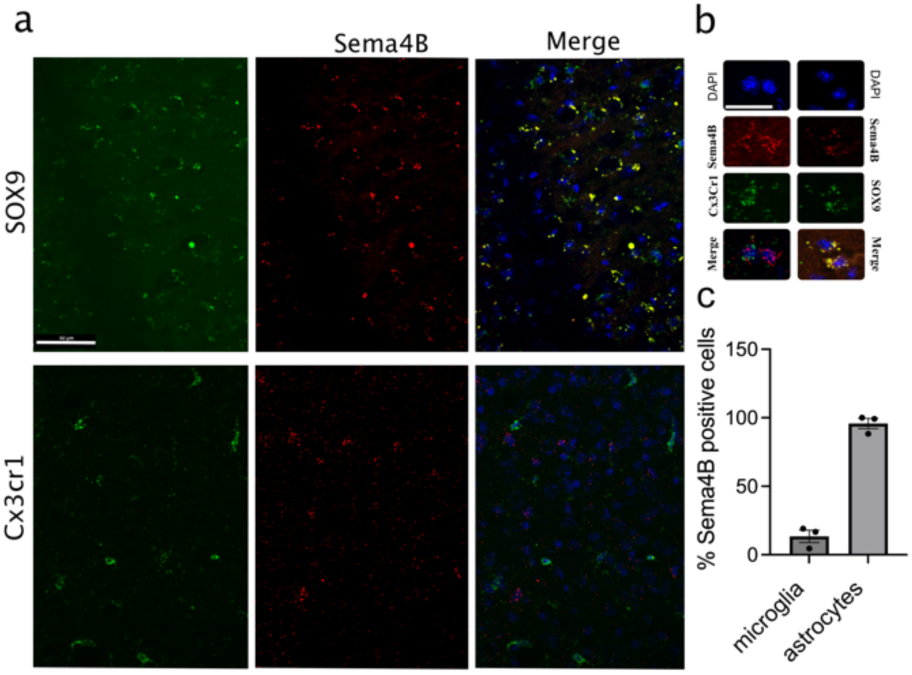
*In situ* RNA expression of Sema4B in the adult cortex. a) Representative cortical images stained with probes for Sema4B and Sox9 (upper panel) or Sema4B and Cx3cr1 (lower panel). Scale bar 50μm. b) Example of microglia and astrocytes expressing Sema4B. Scale bar 25μm. c) The percentage of cells positive for Sema4B/Cx3cr1 and Sema4B/Sox9 is presented (n=3 mice, each dot is an average of 3 sections per mouse).

### The inflammatory response is altered in Sema4B knockout mice

The mRNA expression of various cytokines was examined to test the inflammatory response in Sema4B knockout mice following injury, (Fig 2a; S1a, b). In Sema4B knockout (Sema4B^−/−^) mice, some proinflammatory factors (CCL2, CCL3, CCL5, and IL6) were significantly lower 12h after injury. However, at 18h, CCL2 and IL6 were more highly expressed in the Sema4B knockout mice, suggesting the differences may represent a shift in the kinetics. At the protein level, we tested IL-12, IL-18, IL10, and IL-1β 24 hours after injury using ELISA (Fig 2b; S1c, d). Both IL-12 and IL-1β were lower in Sema4B^−/−^ 24h after injury, although only IL-12 reached statistical significance.

**Figure 2:**
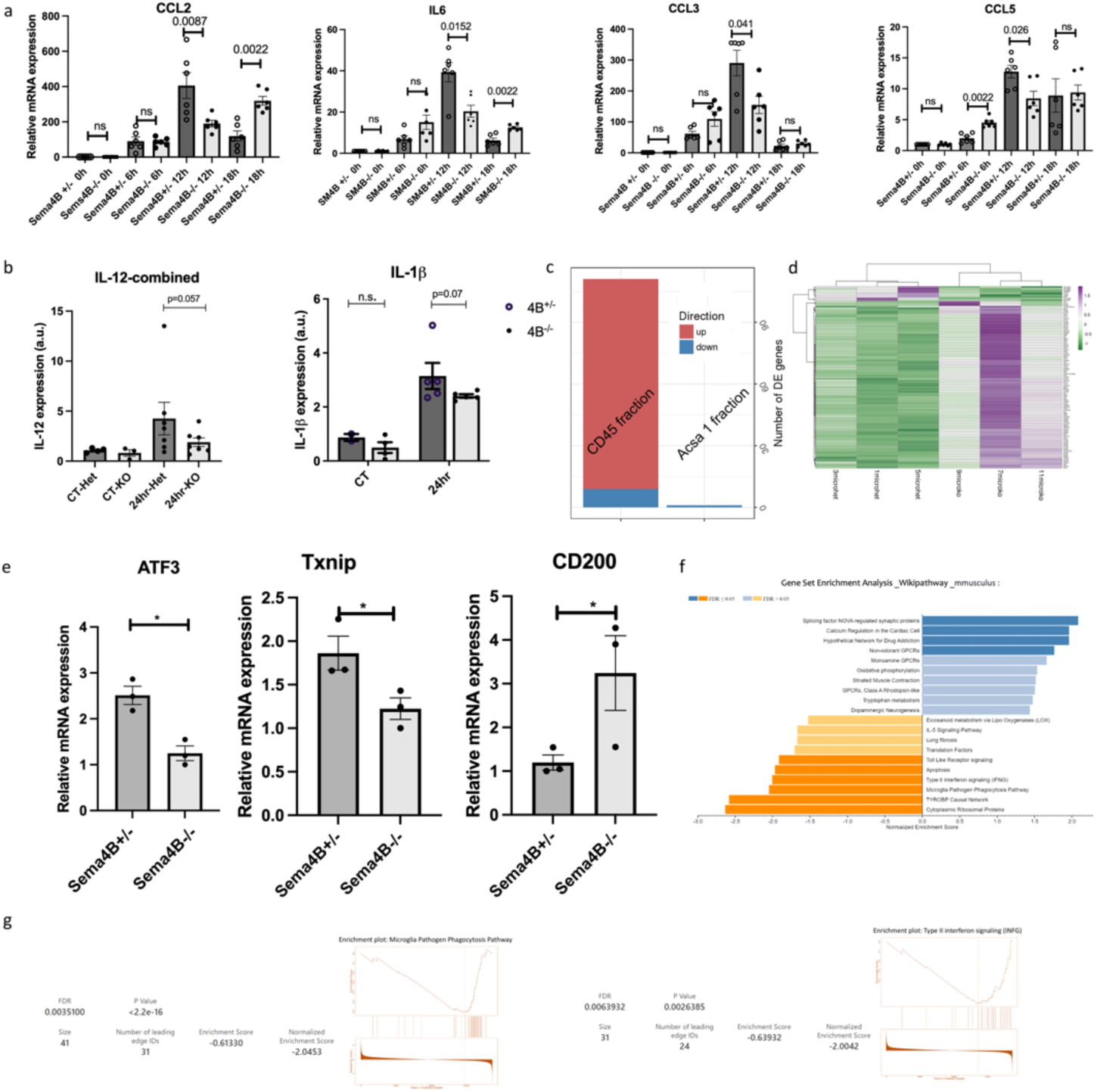
The neuroinflammatory response following cortical stab wound injury is attenuated in the absence of Sema4B. a. qPCR analysis of the cortical tissue near the site of injury isolated from Sema4B+/− and Sema4B-/- mice at the indicated time points after stab injury (n=6; p<0.05; two tail Mann Whitney U test). b. Levels of proinflammatory cytokines 24h after cortical injury at the site of injury were evaluated using ELISA (n=5-7; p=0.02; Fisher’s combined probability test). c. RNAseq analysis of the immune and astrocytic cell fractions isolated by panning 12h after cortical injury. Plots summarizing the overall number of differentially expressed genes (DEGs). Genes with Padj < 0.05 and |log2FoldChange (FC)| > 1 were identified as significant DEGs. (n=3 repeats, with 5 mice per genotype within each repeat). d. Heatmap displaying normalized expression levels of significantly dysregulated genes (I LogFC I > 1; p-adj value < 0.05) in microglial fraction. e. qPCR analysis of the immune cell fraction isolated by panning 12h after injury (n=3; p=0.05; Mann Whitney one-tailed test). f. Gene Set Enrichment Analysis (GSEA) showing significantly dysregulated canonical pathways in astrocytic Sema4B Knockout microglia. GSEA was performed using the Wikipathway database (Mus musculus) with genes pre-ranked according to their log2Fold change. Up-regulated gene-sets are highlighted in blue and downregulated pathways in orange. g. Enrichment plots for two of the most downregulated pathways in knockout microglia/macrophages: microglia pathogen phagocytosis pathway (left) and type-II interferon signaling (right).

To test the changes in gene expression more systematically, we isolated astrocytes and immune cells from the injury site 12h after the injury and performed RNAseq. When comparing the astrocytes from Sema4b+/− and Sema4B-/- mice, we detected only one differentially expressed gene (GM10800, Fig 2c). In contrast, 102 upregulated and 9 downregulated genes in the immune fraction of Sema4B-/- mice were identified (Fig 2d). We validated these results by repeating the immunopanning separation and testing 3 genes from the list in the immune cell fraction (Fig 2e). Since inflammation in Sema4B-/- mice is altered, we focused on genes related to the immune response. To find these genes, we used the gene set enrichment analysis (SEGA) software using ‘explore the molecular signatures’ database (MSigDB) (13, 14). We tested ATF3, a negative regulator of the neuroinflammatory response, and as such might be expected to dampen the inflammatory response of microglia/macrophages (15). Txnip mediates glucocorticoid-activated NLRP3 inflammatory signaling in mouse microglia and was also found to interact with nucleotide-binding oligomerization domain-like receptor protein 3 (NLRP3), which activates the NLRP3 inflammasome and promotes inflammatory processes (16). Finally, we also tested CD200, since this signaling plays a significant role in maintaining microglia in a quiescent state and functions as an anti-inflammatory signal (17). Although CD200 is mostly expressed by neurons, its expression is upregulated in microglia following Kainic acid injection (18). To get a sense of the changes in gene expression in the Sema4B knockout we used the list of the genes pre-ranked according to their log2Fold change. We then used the Wikipathway database (Mus musculus) to map the pre-ranked genes to their respective enrichment pathways and gene sets (Fig 2f). The GSCA visualization of the gene sets in a few pathways is presented (fig S2) and a heat map of the leading edge of 2 pathways related to microglia functions including toll-like receptor microglia phagocytosis and INFG signaling are also presented (Fig 2g).

### Altered response of microglia/macrophages in the absence of Sema4B

To test microglia/macrophage reactivity, we used a morphology analysis using IbaI to visualize the cells. The cell morphology changes from a ramified state to a more amoeboid shape as it becomes more reactive (Fig 3a). The cells were stained for Iba1 24h after injury, and the morphological analysis was performed (Fig 3a-c). The results are consistent with enhanced microglia/macrophage reactivity in Sema4B heterozygous mice as compared to Sema4B knockout mice, though the number of Iba1-positive cells is the same (Fig 3da-d). Importantly, there were no morphological differences between genotypes in non-injured mice, indicating that there is no difference in their basal activation state (Fig 3a-g). To get a sense of the significance of the morphological analysis between the two genotypes we challenged wild-type mice with LPS and tested their morphology after 24h. The differences with and without LPS were small and did not reach statistical significance (Fig S3). We, therefore, conclude the differences detected between Sema4B knockouts and heterozygous after injury are biologically meaningful.

**Figure 3:**
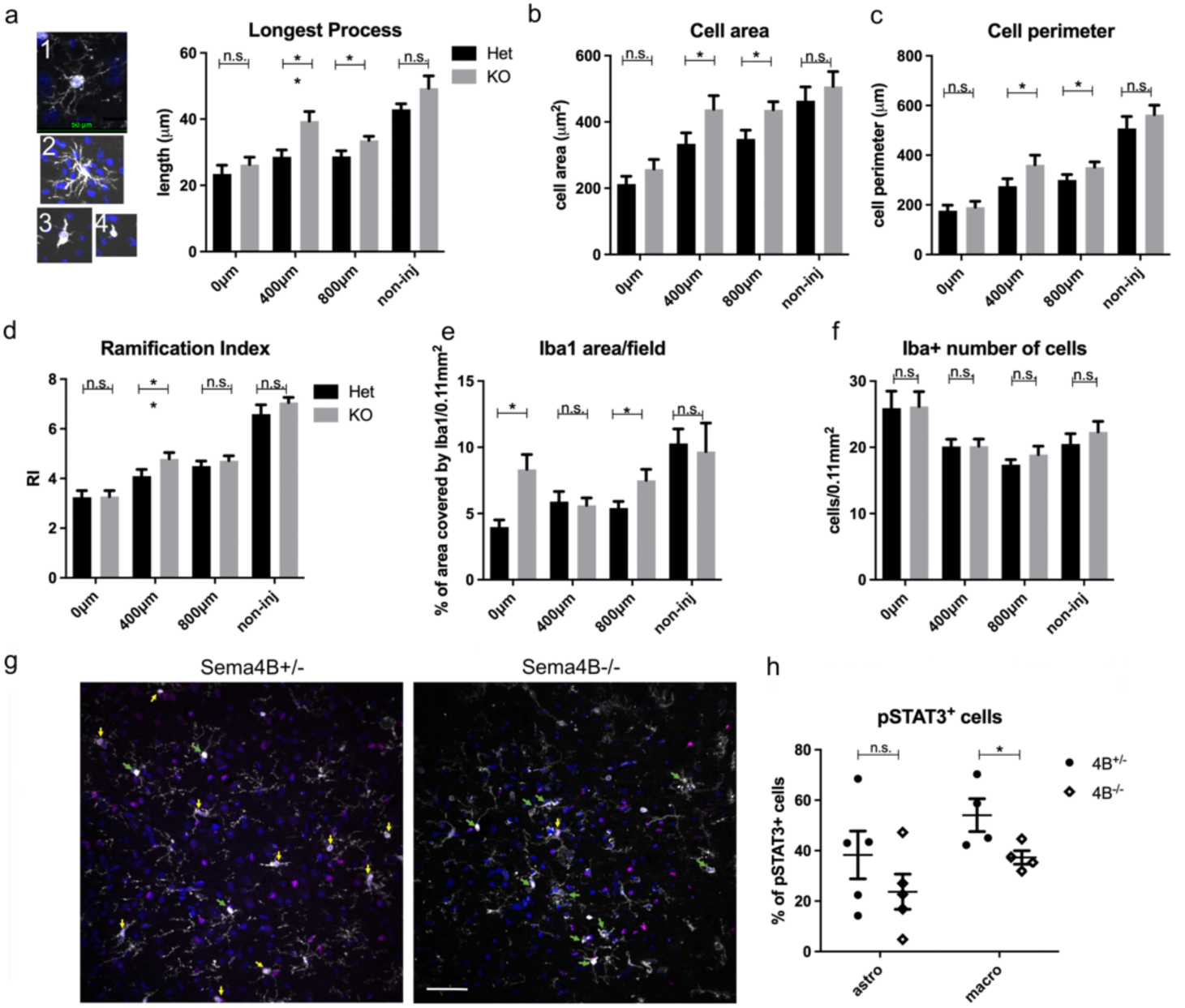
Morphology of microglia/macrophages in Sema4B mutant mice is altered and reactivity is attenuated. a) examples of microglia cells stained with IbaI from most ramified (1) to the most ameboid cells (4). b-e) Morphological measurements of Iba1 images in Sema4B^+/−^ and Sema4B^−/−^ mice including the longest process (a), cell area (b), cell perimeter (c), ramification index (d), and Iba1 area/field (n=3-4 mice, 3 sections/mouse; Mann-Whitney one-tailed test). f) Quantification of the density of Iba1 positive cells (n=5 mice; Mann-Whitney one-tailed test). g) Representative images of Iba1 (gray) and pSTAT3 (magenta) 24h after cortical injury. Green arrows mark pSTAT3 positive/Iba1 negative, and yellow arrows mark pSTAT3positive/Iba1positive cells (Scale bar 50μm). h) Quantification of Iba1/pSTAT3 and S100b/pSTAT3 cells in Sema4B^+/−^ and Sema4B^−/−^ 24h after injury (n=4-5 mice, 3 sections per mouse; Mann-Whitney one-tailed test).

Following injury, microglia change from a homeostatic state into a reactive state, a process that involves the downregulation of homeostatic genes. To monitor the homeostatic gene response following injury we tested Hexb, Tgfb1, p2yr12, p2yr13, and Tmem119 at 0h, 6h, 12h, and 24h after injury several genes in wild-type mice (Fig S4). Overall, all homeostatic genes were downregulated after injury (Fig S4).

An additional aspect of immune activation after injury manifests in the migration of immune cells toward the site of injury (19). To further study the impact of Sema4B, we tested the cell density of Tmem119 (a more microglia-specific marker), CD45 positive cells (general immune cell marker), F4/80+ (macrophage marker), F4/80+/Tmem119-, F4/80+/Tmem119+ cell density, and the % of Tmem119+ cells out of the total Iba1 cells (Fig S5). Together, these results indicate that there is no difference in microglial/immune cell migration/accumulation in the injury site in Sema4B mutant mice.

INFg signaling found to be repressed in Sema4B knockout mice (fig. 2f, g) is primarily mediated by Stat signaling. We therefore tested Stat3, a major transcriptional activator involved in the injury response of microglia/macrophages and astrocytes. Without injury, pSTAT3 is undetectable in the mouse cortex (Fig. S6a). To test the state of pSTAT3 after injury we stained for Iba1 and pSTAT3 (microglia/macrophages, Fig 3j, S6b), and S100b and pSTAT3 (astrocytes, Fig S6c). We detected a significant reduction in the level of pSTAT3 in the microglia/macrophage population, but not in astrocytes (Fig 3k).

### Astrocytic Sema4B modulates the reactivity of microglial cells *in vitro*

To assess whether Sema4B can act directly on microglia, we tested if Sema4B has a binding site on these cells and found that they do indeed (Fig 4a). To test the potential effect of Sema4B on microglia, we used an *in vitro* system of injury. Conditioned medium was collected from injured mixed glial cells (we named this “injury supernatant” or “iSup”). We tested the expression of different proinflammatory cytokines (CCL2, IL6, CXCL10, IL1α, IL1β, and TNF) after microglial stimulation with iSup with or without Sema4B for 3h (Fig 4b, and Fig S7a-c) or 9h (Fig 4c, and Fig S7d-f). Sema4B by itself was a weak stimulus of microglia in the case of some genes. However, together with iSup, Sema4B activated some proinflammatory signals more robustly than just iSup alone. To further test the impact of Sema4B on microglia, we also tested the percentage of microglia expressing iNOS 24hr after stimulation with iSup and Sema4B (Fig 4d, and Fig S8). Consistent with the qPCR results, we detected that more microglial cells express iNOS following exposure to Sema4B and iSup. Finally, to test if astrocytic Sema4B is indeed a modulator of microglial cell activation, we incubated wild-type microglia with Sema4B heterozygous and knockout astrocytes. The culture was stimulated with iSup and the expression of a few inflammatory markers (Chil3, Psmb8, CD300fl) that are either expressed predominantly by microglia (Chil3, CD300fl) or show differential expression only in cultures of both astrocytes and microglia (Psmb8) were tested (Fig 4f). Our results clearly show that astrocytic Sema4B is a modulator of microglia, while the limited expression of Sema4B on microglia is not enough to compensate for it.

**Figure 4:**
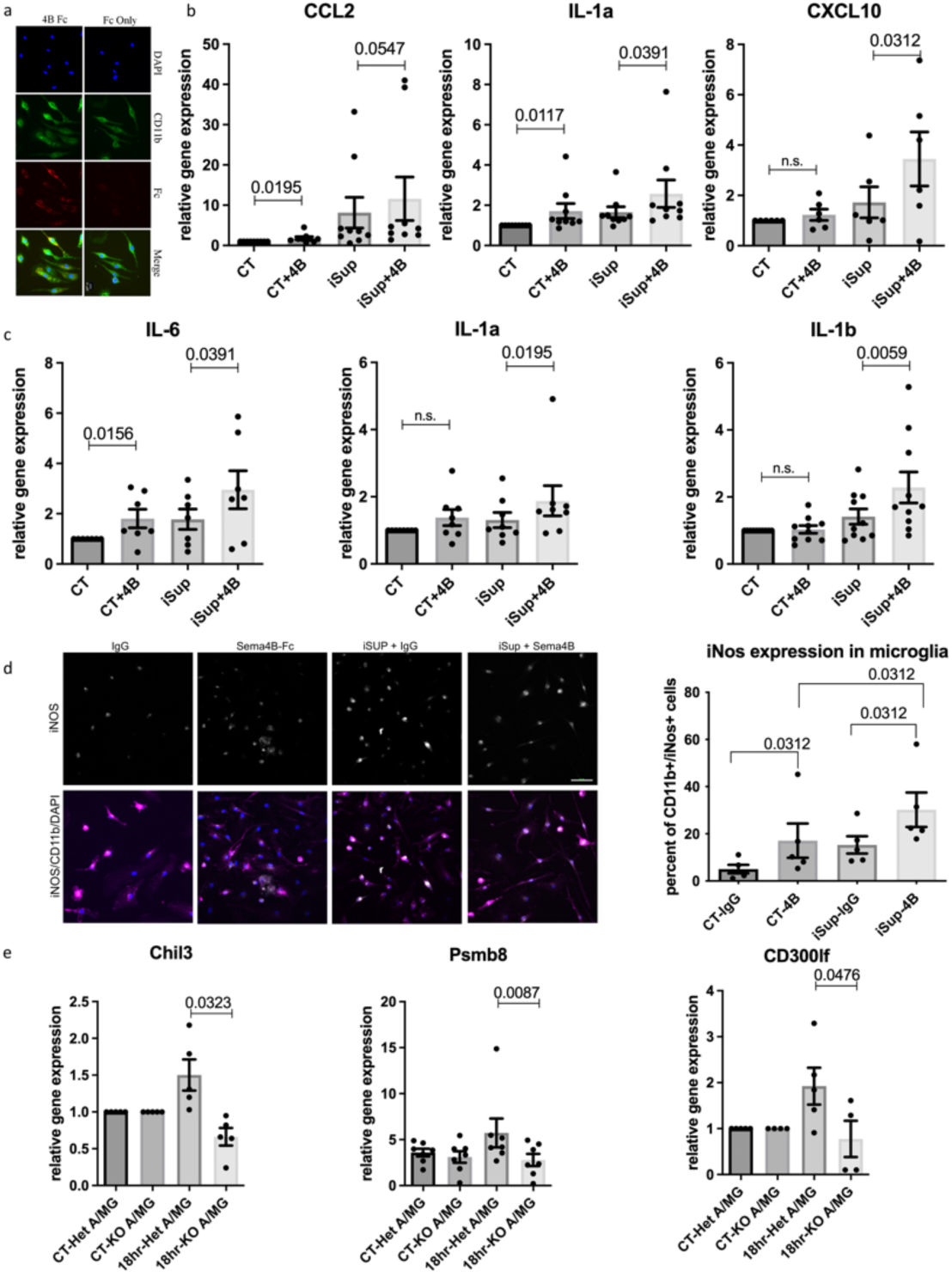
Sema4B amplifies the reactivity of cultured microglia following injury. a) Live staining of cortical cultured microglia with Sema4B-Fc demonstrates a binding site for this ligand. (Scale bar 10μm). b) qPCR analysis of cultured microglia incubated with HEK293 cells expressing the full length Sema4B or GFP, and/or iSup for 3h (n=6-9 experiments; p<0.05; Wilcoxon one-tailed test). c) qPCR analysis of cultured microglia treated with Sema4B or GFP, and/or iSup for 9h (n=7-10 experiments; p<0.05; Wilcoxon one-tailed test). d) Representative images of iNOS (gray) and CD11b (magenta) staining of microglial cultured cells expressing iNOS 24h after treatment (Scale bar 50μm). Quantification of microglial cultured cells expressing iNOS 24h after treatment with either an IgG control or Sema4B-Fc and/or iSup from all images (n=4 experiments, 3 fields/experiment; Wilcoxon one-tailed test). e) qPCR analysis of co-cultured wild-type microglia with astrocytes isolated either from Sema4B^+/−^or Sema4B^−/−^ mice and treated with iSup for 18h (n=6 experiments, Wilcoxon one-tailed test).

### Plexin-B2 is the receptor of Sema4B in microglia

The Sema4B receptor in microglia is unknown. The likely candidate is Plexin-B2, which was suggested to be a receptor based on a pErbB2 phosphorylation assay (20). We used qPCR to examine the expression of Plexin-B2 and its related family member, Plexin-B1, in cortical microglia and in the BV2 microglial cell line. In both, Plexin-B2 is highly expressed while Plexin-B1 is expressed at much lower levels (Fig 5a). To test if Sema4B can activate PlexinB2, we used a COS cell collapse assay. Indeed, Sema4B can activate this receptor and cause cell collapse (Fig 5b, c). To investigate the potential role of Plexin-B2 in microglial activation following injury, we crossed Cx3cr1CreER (21). Cells targeted with this system include microglia, non-parenchymal CNS macrophages, and selected peripheral macrophages, but notably, exclude shorter-lived monocytes. The protein expression of Plexin-B2 with or without 4-hydroxytamoxifen (4OHT) was tested in microglia i*n vitro*. As expected, Plexin-B2 is expressed by microglia in culture and this expression is eliminated after the 4OHT treatment (Fig S9a). In response to iSup activation, microglia expressed higher levels of cytokines. This response, however, was no longer enhanced by Sema4B after 4OHT treatment even in control microglia. In fact, it had the opposite effect (Fig S8b). As an alternative, we used the microglial cell line BV2 to test the function of Plexin-B2. Targeting Plexin-B2 with shRNA reduced Plexin-B2 expression both at the RNA and protein levels (Fig 5d, e). We tested the expression of a few cytokines in BV2 cells after stimulation with Sema4B (Fig f). Sema4B by itself triggered expression (CXCL10, TNFα, CCL2) while cells with a Plexin-B2 knockdown did not respond to Sema4B. In most cases, the activation of BV2 by iSup was not additive to Sema4B (not shown). In contrast, IL-1b expression was not affected by Sema4B alone but was higher with iSup together with Sema4B. In this case, shRNAs targeting PlexinB2 resulted in the loss of the additive impact of Sema4B and iSup (Fig 5f).

**Figure 5:**
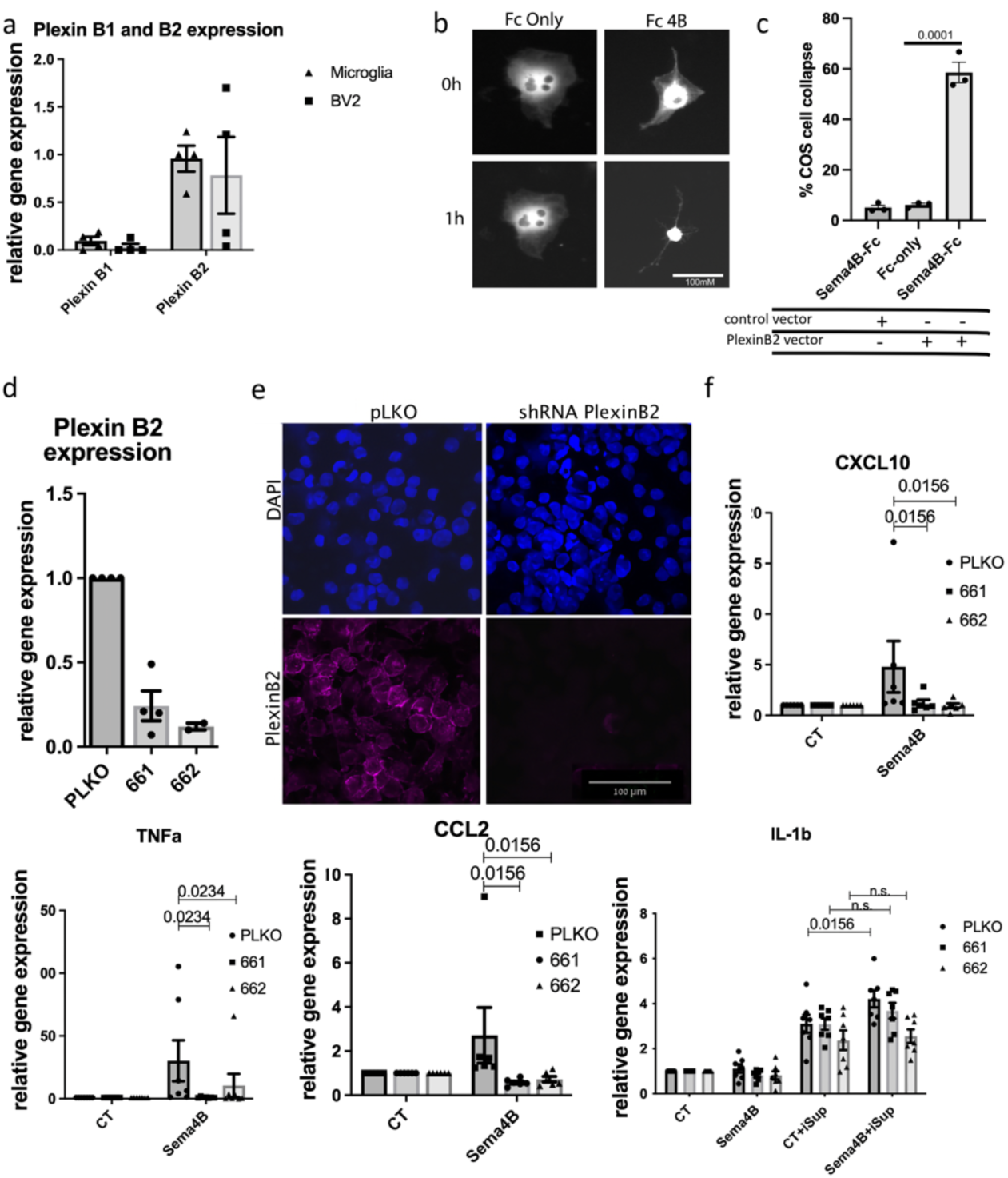
Plexin-B2 is the likely receptor for Sema4B in microglia. a) qPCR analysis of Plexin-B1 and B2 expression in cultured microglia and BV2 microglial cell line (n=3). b-c) COS cell collapse assay was used to test the potential of Sema4B to function as a ligand for Plexin-B2. b) representative image of a COS cell before and 1h after treatment with Sema4B-Fc or Fc only. (Scale bar 100μm). c) Quantification of COS cell collapse is shown (n=3, t-test). d) qPCR analysis of Plexin-B2 expression in BV2 cells treated with two shRNA sequences targeting Plexin-B2 (n=3 experiments). e) Representative image of Plexin-B2 immunofluorescence of BV2 cells with scrambled shRNA sequence or shRNA targeting Plexin-B2. f) qPCR was used to analyze the expression of different cytokines (CXCL10, TNFα, CCL2, IL1β) in BV2 cells pretreated with Plexin-B2 shRNA. The BV2 cells were then co-cultured with HEK293 cells expressing either full-length Sema4B or GFP, in the presence or absence of iSup for 3hrs (n=6-7 experiments; Wilcoxon one-tailed test).

### Microglial Plexin-B2 regulates microglial activation after cortical injury

To test if Plexin-B2 is important for microglial activation, we used the Cx3cr1creER:PlexinB2^fl/fl^ mouse model to perform cortical injury. We used in situ hybridization for PlexinB2 and CX3Cr1 (microglia/macrophage marker) on PlexinB2^+/fl^ and PlexinB2^fl/fl^. Analysis of Plexin-B2 expression in Cx3cr1creER:PlexinB2^fl/fl^ mice 3 weeks post-TAM treatment and 3 days after injury confirmed Plexin-B2 expression is almost exclusively detected in microglia/macrophage cells and that there is almost no expression in knockout cells (Fig 6a). To test if the absence of Plexin-B2 in microglia/macrophages attenuates activation after injury, we performed a morphological analysis 24hr after injury (Fig 6b-d). Interestingly, microglial/macrophage reactivity is attenuated, although less than what we observed in Sema4B knockout mice. Since in the absence of Sema4B, neurons are more protected after injury (12), we tested if this is the case in Cx3cr1creER:PlexinB2^fl/fl^ mice as well. For this, we stained tissues using slices with FJC 24h after injury. Indeed, as in the case of Sema4B, fewer neurons died in the absence of PlexinB2 in microglia/macrophages.

**Figure 6:**
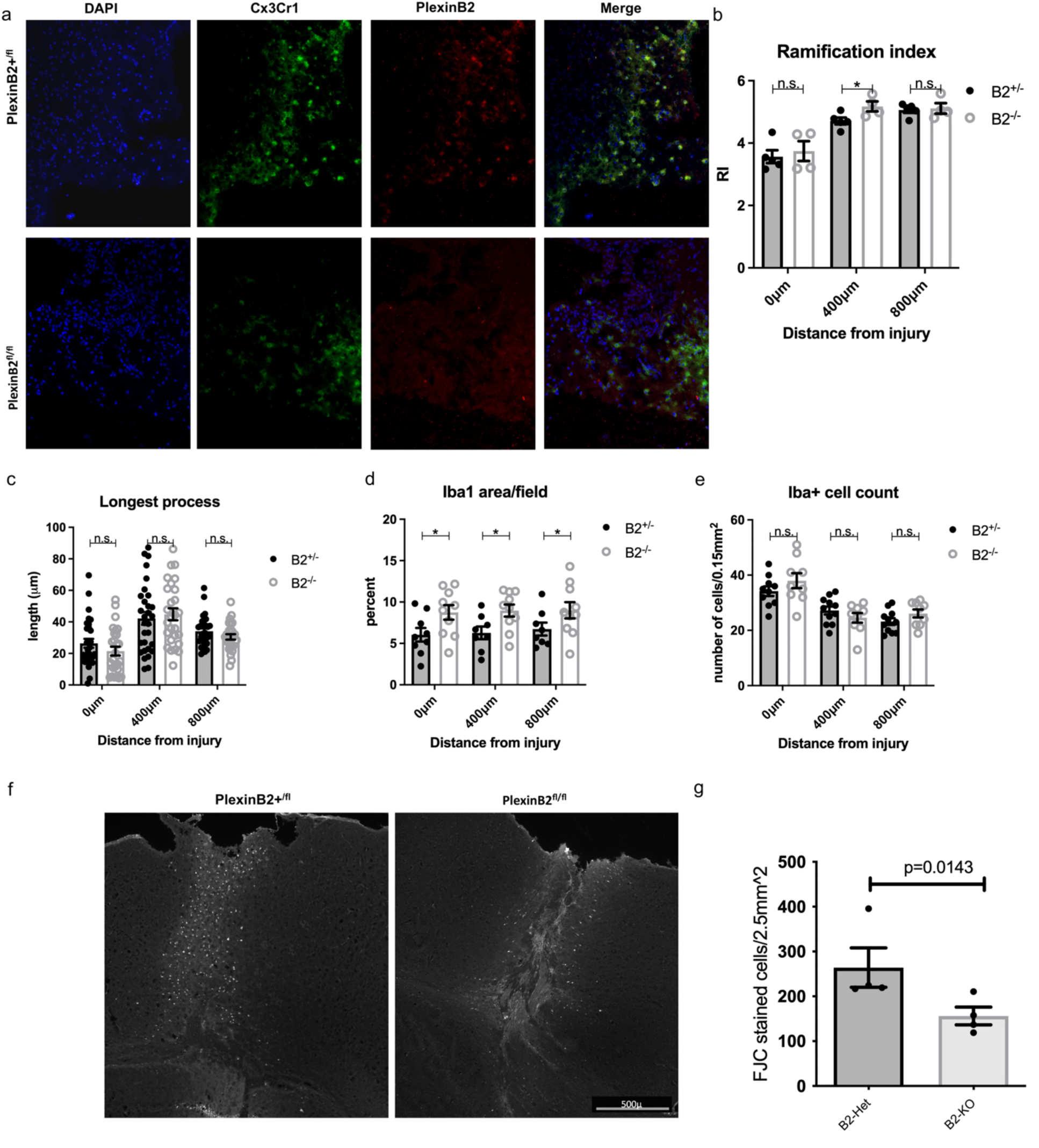
Microglia/macrophage activation is attenuated in Cx3cr1creER:PlexinB2^fl/fl^ mice. a) In situ hybridization of Plexin-B2 and Cx3Cr1 on Cx3cr1creER:PlexinB2^+/fl^ and Cx3cr1creER:PlexinB2^fl/fl^ mice, 3 weeks post-TAM treatment and 3 days after injury. note that most cells expressing PlexinB2 are Cx3Cr1 positive, and that this expression is almost undetected in PlexinB2^fl/fl^ mice. b-e) Morphological measurements of Iba1 images in Cx3cr1creER:PlexinB2^fl/fl^ and PlexinB2^+/fl^ control mice (n=4-5, 3 sections/mouse; Mann-Whitney one-tailed test). f) Representative coronal cortical sections in Cx3cr1creER:PlexinB2^fl/fl^ and PlexinB2^+/fl^ mice 1d post injury. The panel shows dead neurons labeled with FJC. g. Quantification of the total number of dead neurons (FJC positive) 1d post-injury (n=4 mice, 4-6 sections/mouse; p=0.014; Mann-Whitney one-tailed test).

Finally, to test if Plexin-B2 in microglia and Sema4B are part of the same genetic cascade, we analyzed the reactivity of microglia/macrophages in double heterozygotes for both microglial Plexin-B2 and Sema4B. We detected some changes in microglia morphology consistent with a reduction in microglial reactivity after injury in these double heterozygous mice (Fig 7 a-f), supporting a link between Plexin-B2 on microglia and astrocytic Sema4B in the same genetic signaling cascade.

**Figure 7:**
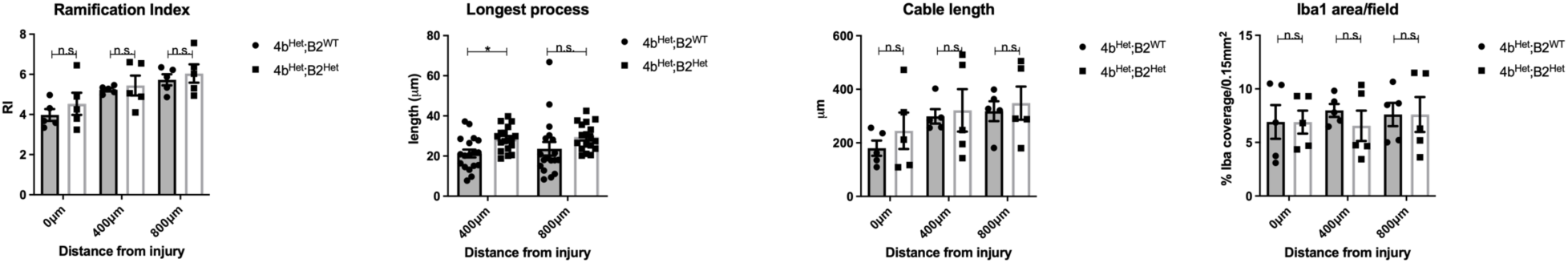
*Morphological measurements of Iba1 images in* Cx3cr1creER:PlexinB2 mutant and control mice (n=5; Mann-Whitney one-tailed test). Morphological measurements of Iba1 images in Cx3cr1creER:PlexinB2 mutant and control mice (n=5, 3 sections/mouse; Mann-Whitney one-tailed test).

## Discussion

Brain injury triggers a response aimed at restoring homeostasis and limiting damage. A critical part of this response requires coordinated activation of microglia/macrophages and astrocytes. Here we showed that following cortical injury, Sema4B enhances the reactivity of microglia/macrophages via the Plexin-B2 receptor, and thus, the crosstalk between Sema4B and Plexin-B2 modulates the microglial/macrophage proinflammatory response following injury.

### Sema4B is a modulator of inflammatory response

Semaphorins were originally identified as axon-guidance molecules. Over the years, other functions have been discovered, including in the immune system (22). Many semaphorins with immunological function belong to the type 4 semaphorins and include Sema4D (also named CD100) as a prominent example (22). In the CNS, Sema4D was shown to modulate microglial cells. For instance, in culture, Sema4D enhances CD40-mediated activation of microglial cells (23). Consistent with this, blocking antibodies for Sema4D reduces the activation of microglial cells in LPC-induced lesions (24). Moreover, in ischemic stroke, activation of microglia is attenuated, and cell death is reduced in Sema4D knockout mice (25). It seems that Sema4B may play a similar role. First, we have recently shown that in cortical injury neuronal cell death is reduced in the absence of Sema4B (12). Moreover, here we show that in the absence of Sema4B in mice, activation of microglial cells attenuated. Consistent with this, cultured microglia challenged by injury are more highly activated in the presence of Sema4B. Therefore, both Sema4B and Sema4D under conditions of brain injury act as ligands to promote a pro-inflammatory response.

### Plexin-B2 is a microglial receptor for Sema4B

Plexin-B1 and B2 are considered type 4 semaphorin receptors, although Sema4D can also act via CD72 in certain immune cell functions (10). In microglia, both RNA sequencing and single-cell RNA sequencing studies suggest that Plexin-B2 is the primary PlexinB-expressed gene, whereas Plexin-B1 is highly expressed in astrocytes, as reported before (26–28)(26–28) Moreover, recent findings show that Plexin-B2 plays a crucial role as a receptor in microglia/macrophages following spinal cord injury, regulating cell corralling, and that its absence can lead to diffuse tissue damage, inflammatory spillover, and hindered axon regeneration (29).

The Sema4B receptor is still incompletely defined, although Plexin-B2 mutants and ErbB2 phosphorylation assays strongly support Plexin-B2 as the Sema4B receptor in epithelial cells (20). Our own experiments show that Sema4B can bind Plexin-B2 and serve as a functional receptor in a COS cell collapse assay. We also find in a microglial cell line that Sema4B-induced cytokine expression is Plexin-B2 dependent. Moreover, the similar impact of Sema4B and Plexin-B2 knockouts on microglial reactivity is consistent with crosstalk between these two –receptor-ligand pairs, a conclusion supported by the genetic interaction between these two genes. However, these findings appear to be at odds with earlier research showing that Plexin-B1 is the microglial receptor mediating Sema4D activity (23). One possible explanation for this discrepancy is the difference between cortical microglia used here (as well as in the studies of (28) and (26)) and the use of microglia extracted from the whole brain used by Okuno et al. (23).

### Does Sema4B function as a ligand or receptor after brain injury?

Our earlier study showed that 7 days after injury, astrocyte proliferation is diminished while activation is changed, and our *in vitro* experiments indicate that injury-induced astrocyte proliferation is cell-autonomous and reliant on Sema4B phosphorylation, thus suggesting Sema4B’s role as a receptor in astrocytes (11). In contrast, the impact of Sema4B on neuroinflammation in general, and microglia specifically, presented in the current study shows a ligand function of Sema4B. Interestingly, although both Plexin-B2 deletion in microglia and Sema4B knockout affect microglial reactivity, the latter seems to have a more pronounced impact. This may be explained either by the reduced efficacy of the tamoxifen-induced Plexin-B2 deletion or by a dual function of Sema4B both as a ligand and as a receptor. Future studies will be needed to differentiate between these possibilities.

### Microglia-astrocyte communication via Plexin-B2 and Sema4B

The findings presented in this study, combined with earlier work, show that Sema4B is primarily expressed by cortical astrocytes, with some expression seen in a minority subpopulation of microglia (11, 27, 30), and that astrocytic Sema4B is necessary for the modulation of microglial reactivity after injury in an astrocyte-microglia co-culture system. Based on these results, we propose a model in which under conditions of injury, astrocytic Sema4B acts through the Plexin-B2 receptor to enhance DAMP-triggered activation of microglia. As a result, neuroinflammation following a cortical stab wound injury is amplified.

In summary, our study provides novel insights into the complex cellular mechanisms involved in the response to brain injury and highlights the potential of Sema4B and Plexin-B2 as a therapeutic target for intervention.

## Materials and methods

### Antibody and primer information

can be found in the supplementary methods.

### Animals and surgical procedure

Animal handling adhered strictly to national and institutional guidelines for animal research and was approved by the Ethics Committee of the Hebrew University.

Sema4B+/− mutant mice were bought from the Mutant Mouse Regional Resource Center (MMRRC) and used as described previously (11). The generation of PlxnB2 conditional knockout mice was described previously (31). CX3Cr1CreER mice from (21) were crossed with floxed Plexin B2 mice for the KO experiments. C57BL/6 wild-type mice were obtained from ENVIGO (Rehovot, Israel). All cortical injury experiments were performed on mice aged 7−12 weeks old. Genotype was determined by PCR analysis of genomic DNA isolated from ear clippings of 3-week-old mice (additional information can be found in the supplementary). In all experiments, both males and females were used in similar numbers except for Fig S4 in which we used males only. Extended information regarding the mice work, Tamoxifen treatment, and Stab wound injury model is in the supplementary method.

### Cell Culture

Astrocytes and microglia were cultured and purified as described in the supplementary methods.

### Immunofluorescence, Fluoro-Jade C staining, RNA extraction and qPCR

The explanation is found in the supplementary methods.

### Protein harvesting and ELISA

Protein harvesting explanation is found in the supplementary methods and ELISA was performed according to the manufacture protocol using mouse IL-18 ELISA kit (MBL, #7625, Woburn, MA, USA) and R&D Systems (Minneapolis, MN, USA) mouse IL-10 (#M1000B), mouse IL-12 (#M1240), and mouse IL-1β (#SMLB00C) kits.

### Microglial morphological analysis

Cell analyses were performed using ImageJ software. Tissue sections were prepared from mice 1 day after stab injury, stained with Iba1, and photographed at 40x enlargement in a spinning disc confocal microscope (Nikon, Tokyo, Japan) in increments of 1.5-3mm thick optical sections. The sections were photographed directly next to the injury, 400, & 800µm away from the injury, and in the non-injured cortical hemisphere. In ImageJ, the Z stack was amalgamated using maximum intensity. For the longest cell process, the 3 most ameboid cells per field were analyzed and measured. For the average Iba1 area/field, the upper threshold of each picture was set to the same value for every set, and % of area above the threshold was measured. For ramification index (RI) and cable length, we developed an ImageJ script developed for use with ImageJ was and used it to measure these parameters for each cell/field and calculate the average values per field. The average values of each field were calculated for each mouse. Analyses were performed on 3-4 animals of each genotype, and 6-9 fields per animal. The ramification index was calculated using the formula RI=(perimeter/area)/[2(π/area)1/2] as described previously (32) and cable length was defined as the sum of lengths of a cell’s skeleton ramifications.

### RNA seq analysis

#### Cell isolation by immunopanning

First, cortical tissue (3mm x 3mm) around the stab wound injury from 3 mice for each genotype was dissected out and the white matter was peeled off. Cell immunopanning was performed essentially as previously described (33). Cortical tissue was dissociated into cells using 200u/ml papain (cat # LS003126, Worthington, Columbus, Ohio, USA) and DNAse followed by trituration. The cells were added to the secondary-only plate, followed by sequential incubation in plates containing antibodies against CD45, Thy1, O4, and finally ACSA-2. RNA was extracted from plates 1+2 (immune fraction) and the ACSA-2 plate (astrocytes), using Direct-zol^TM^ RNA Mini-Prep (Zymo Research) kit. All samples had a 260/280 OD of at least 1.93 (and an average of 2), and a RIN value of at least 8.

The antibodies used in the panning experiments are indicated in the supplementary section.

#### RNA seq and bioinformatics analysis

##### Sequencing

RNA-seq libraries were prepared at the Crown Genomics institute of the Nancy and Stephen Grand Israel National Center for Personalized Medicine, Weizmann Institute of Science. Libraries were prepared using the INCPM-mRNA-seq protocol (a homemade protocol, similar to Truseq by Illumina). Briefly, the polyA fraction (mRNA) was purified from 500 ng of total input RNA followed by fragmentation and the generation of double-stranded cDNA. After Agencourt Ampure XP beads cleanup (Beckman Coulter), end repair, A base addition, adapter ligation and PCR amplification steps were performed. Libraries were quantified by Qubit (Thermo fisher scientific) and TapeStation (Agilent). Sequencing was done on a single lane on a Hiseq2500 V4 50 cycles single read kit (Illumina). The loading concentration was 10 pM, sequencing parametres were: rd1-61, i7-11. Sequencing yielded between 18M and 22M reads per sample (median of 21,630,000 reads). Average quality was above 35 throughout the sequencing cycles. About 95% of the reads aligned to the mouse genome. These included about 15% of multi-aligned reads, 15% of uniquely aligned reads outside of known exons, and ~65% of uniquely aligned reads that matched known exons. Thus, a median of 14,270,000 reads per sample were used for further analysis.

##### Bioinformatics

Poly-A/T stretches, and Illumina adapters were trimmed from the reads using cutadapt (34); resulting reads shorter than 30bp were discarded. Reads were mapped to the *M. musculus* GRCm38 reference genome using STAR (35), supplied with gene annotations downloaded from Ensembl release 92 (STAR was run with EndToEnd option and outFilterMismatchNoverLmax 0.04). Expression levels for each gene were quantified using htseq-count (36). Differentially expressed genes were identified using DESeq2 (37) with the betaPrior, cooksCutoff, and independent filtering parameters set to False. The pipeline was run using snakemake (38). The GEO Submission access number is GSE230069.

##### Gene Set Enrichment Analysis

The analysis was done with all genes pre-ranked according to their log2Fold change. The Wikipathway database (Mus musculus) was employed to map the pre-ranked genes to their respective pathways and gene sets.

##### IPA Pathway analysis

Pathway analysis and enrichment of immune canonical pathways were performed based on the Ingenuity Pathway Analysis (IPA) Interactions Knowledge Base, including direct and indirect relationships between genes and endogenous molecules. Expression data for 1012 genes were included (genes with P value < 0:01). Signaling pathways with z-score > 1.3 were considered significantly dysregulated. Enrichment of immune pathways, macrophage activation, and neuro-inflammation were particularly highlighted.

#### Injury medium (iSup)

iSup was prepared by first washing 10cm plates of cortical mixed cell cultures with cold PBS, and then scraping the cells from the bottom of the plate in 5 ml of cold cell medium with no serum. Cell chunks were triturated first with a 1ml tip, and then a 200ul tip to break them up. The medium was then vortexed and centrifuged at 1,000g for 10 min at 4°C. The medium was filtered through a 0.22µm strainer, and frozen and stored at −80C. iSup is added to cell cultures at a dilution of 1:4 of the regular volume of medium, along with 2μM ATP (Sigma A2383). Cells are washed once with PBS before RNA is extracted.

#### Sema4B-Fc and Fc control

The generation of Sema4B and Fc control is described in the supplementary method. The use of this reagent was in Figures 4a, 4d, and Figures 5b, 5c. In all other experiments (Figure 4b, 4c, and 5f) recombinant Sema4B was added by adding HEK293 cells transfected with the entire sequence of Sema4B as described below.

#### *Stimulation of* microglial with Sema4B and iSup

HEK293 cells were transfected with plasmids encoding either Sema4b or GFP. 24h later 50,000 microglial cells were cultured with 100,000 HEK293 cells, and the next day, the cells were washed with PBS and incubated with a medium containing no serum (DMEM only) for 24hrs. The cells were treated with iSup and 3-9 hours later the RNA was harvested from the cells, and gene expression levels were measured using mouse-specific primers in qPCR.

#### Plexin-B2 knockdown

For the knockdown experiments, we used MISSION® shRNA lentiviral vectors (pLKO.1) which include puromycin selection marker: TRCN0000078856 (661) and TRCN0000078857 (662). For control, we used a PLKO vector with a scrambled shRNA sequence. Additional details are in the supplementary methods.

#### *In situ* hybridization

Specific probes for SOX9, Sema4B, and Cx3Cr1 were purchased from molecular Instruments (Los Angeles, CA, USA) and were used with their HCR™ RNA-FISH Technology kit according to manufacture protocol (39).

#### COS cells collapse assay

The explanation is found in the supplementary methods.

### Statistical analysis

Values presented are mean ± s.e.m. P-value ≤ 0.05 was considered significant.

Statistical analysis was performed using the two-tailed Mann–Whitney test. Where relevant, P-values were adjusted for multiple comparisons in accordance with the Bonferroni procedure; overall P-values for the different injury experiments were then computed from these adjusted P-values using Fisher’s chi-square test for combined probabilities. For Elisa, we used Fisher’s combined probability test. For microglial activation in culture, we used Wilcoxon one-tailed test.

## Acknowledgments

We thank Dr. Rohini Kuner for providing the PlexinB2^fl/fl^ mice. We thank Thomas Worzfeld and Dr. Luca Tamagnone for their critical reading. We also thank Dr. Luca Tamagnone for providing the PlexinB1 and B2 expression vectors. We also thank the Hebrew University, Faculty of Medicine Core Research Facility, and particularly Drs. Zakhariya Manevitch and Yael Feinstein-Rotkopffor for valuable help with microscopy. We are grateful to Dr. Norman Grover (Department of Experimental Medicine, The Hebrew University) for helpful advice regarding the statistical analyses.

## Author Contributions

N.C. conducted all experiments and contributed to experimental design, manuscript editing, and data analysis for all sections. V.B. conducted most of the experiments in Figure 2 and contributed to the data analysis. A.E and R.E conducted the bioinformatics analysis. R.E. also helped with manuscript editing. O.B. designed all experiments, wrote the manuscript, and supervised all aspects of the work

