## Supplementary figure legends and methods for "Astrocyte-to-microglia communication via Sema4B-Plexin-B2 modulates injury-induced reactivity of microglia"

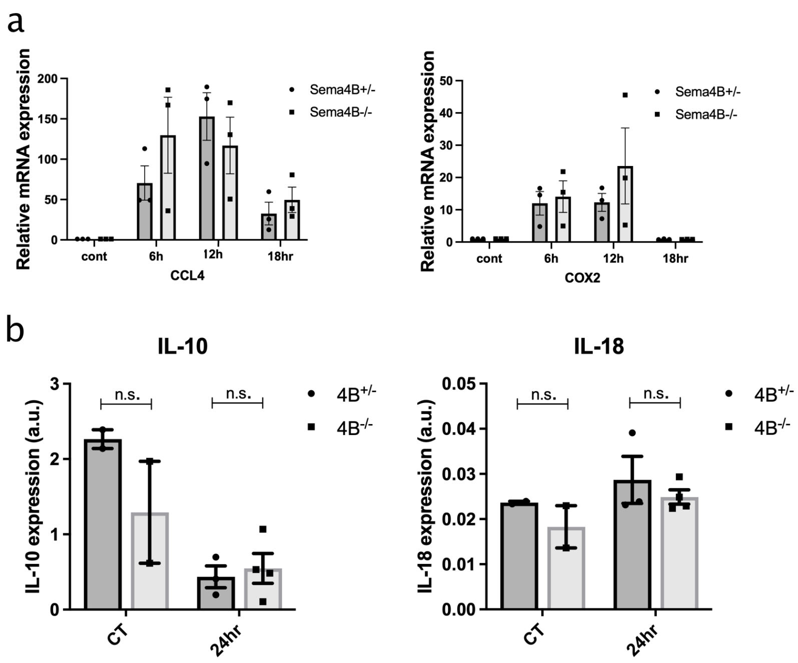


**Figure S1: The inflammatory response following cortical stab wound injury in the absence of Sema4B**

*a) qPCR analysis of the cortical tissue near the site of injury isolated from Sema4B-/- and Sema4B+/- mice at the indicated time points after stab injury (n=4).*

*b) Levels of IL-10 and IL-18, 24h after cortical injury at the site of injury were evaluated using ELISA (n=5-7; Fisher's combined probability test).*

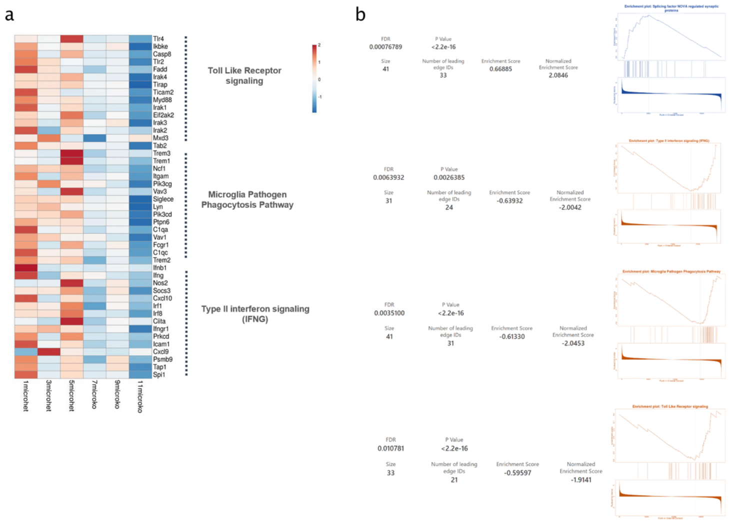


***Figure S2: Pathway Analysis using GSEA and IPA***. *a) Enrichment plots showing significantly dysregulated pathways in microglia from Sema4B-/- mice compared to heterozygous (|NES| > 1.5 and FDR < 0.05). The analysis was performed using the WikiPathways database, and associated statistics were provided alongside each plot. Up-regulated gene sets are highlighted in blue, and downregulated pathways are shown in orange.* ***b)*** *Heatmap displaying the normalized expression of leading genes in three main dysregulated pathways, selected based on GSEA analysis.* ***c)*** *Pathway enrichment using Ingenuity Pathway Analysis (IPA). Bar chart plot displaying the top 50 significant canonical pathways in Sema4B-/- microglia compared to microglia from heterozygous mice (Sema4B+/-). Pathways are ordered by significance (P-value) calculated using a right-tailed Fisher's exact t-test. The color of the bars indicates the enrichment score (Z-score). Orange represents a positive score for upregulated pathways, while blue indicates downregulated pathways with a negative Z-score.*

LPS morphology

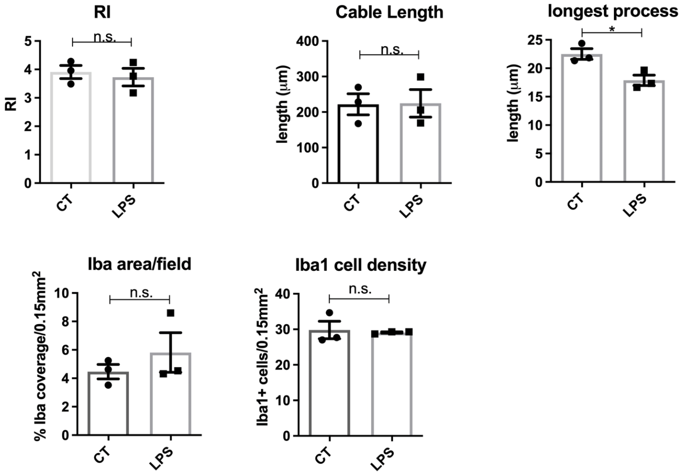


***Figure S3: Morphometric analysis of microglia/macrophage 24h after challenge with LPS***

*Morphological measurements of Iba1 images in Sema4B+/- and Sema4B-/- mice including ramification index, cable length, longest process, Iba1 area/field and quantification of the density of Iba1 positive cells per field (n=3 mice, 3 sections/mouse; Mann-Whitney one-tailed test).*

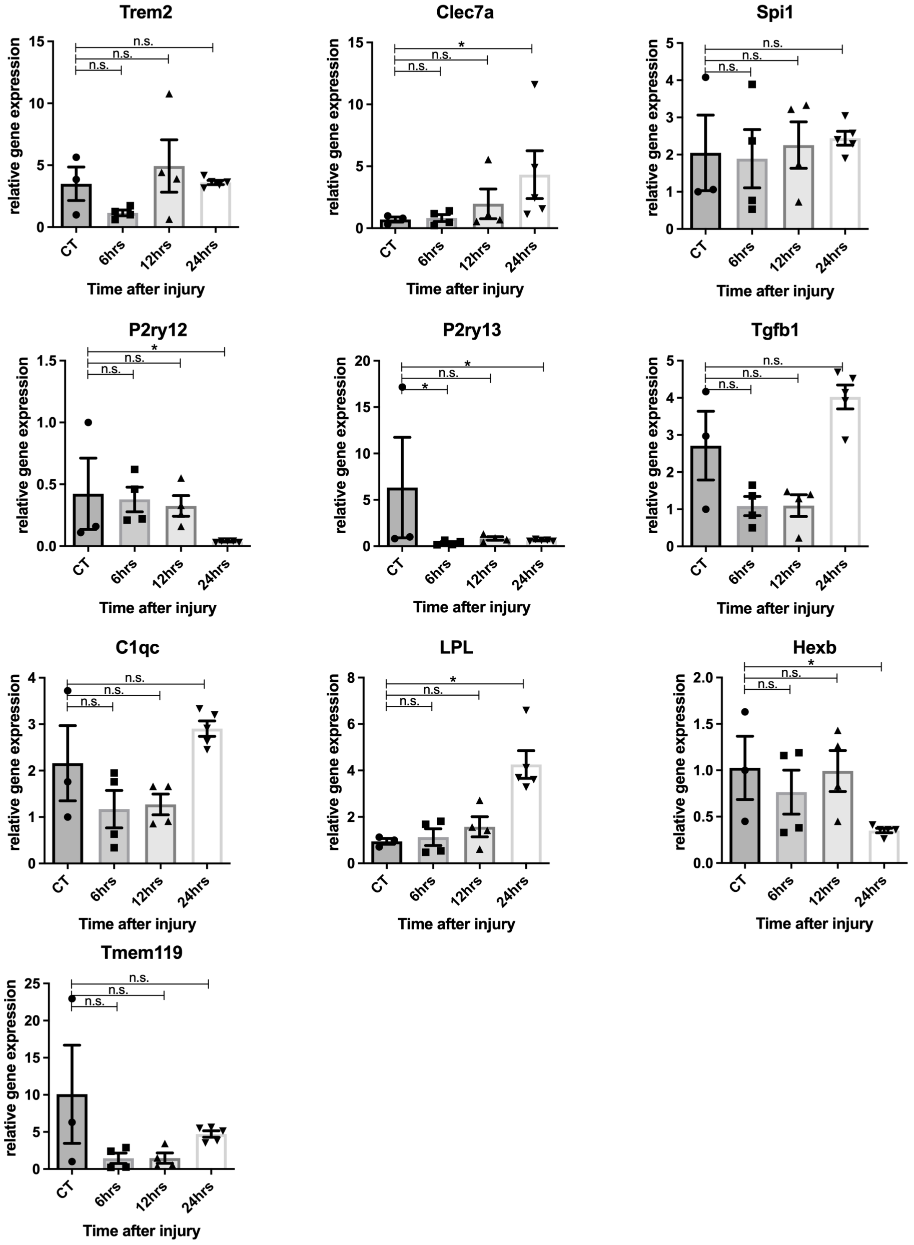
Trem2
Clec7a
Spi1
P2ry12
P2ry13
Tgfb1
C1qc
LPL
Hexb
Tmem119
relative gene expression
Time after injury
CT
6hrs
12hrs
24hrs
n.s.
*

***Figure S4 Expression of reactive and homeostatic microglial markers after injury.***

*qPCR analysis of the cortical tissue near the site of injury isolated from wild-type male mice at the indicated time points after stab injury (n=3-5).*

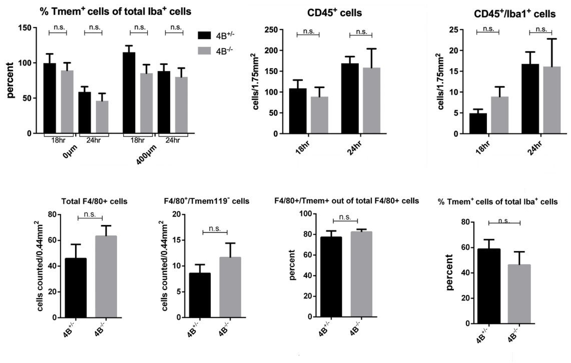


***Figure S5: Immune cell density near the site of injury***

*Quantification of the density of microglia/macrophages and immune cells using Tmem119, CD45, Iba1, and F4/80 positive cells 18h or 24h after injury (n=3-4; Mann-Whitney one-tailed test).*

h) *Quantification* of the density of CD45 positive cells, 18h and 24h after injury is shown (n=4-5 mice, 3 sections/mouse; *Mann-Whitney one-tailed test*).

i) *Quantification* of the density of $CD45^+/Iba1^+$ cells *(n=3-4 mice, 3 sections/mouse; Mann-Whitney one-tailed test)*.

*g) Percent of the Tmem119 positive cells out of the total Iba1 positive cells, 18h and 24h after injury is shown* (n=4-5 mice, 3 sections/mice; *Mann-Whitney one-tailed test*).

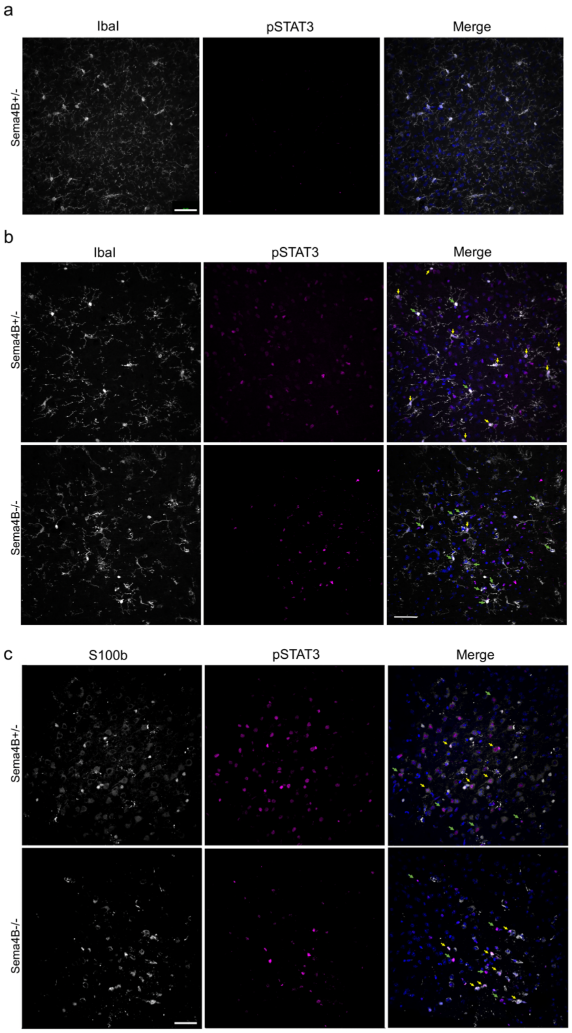


***Figure S6: immunostaining of S100b and pStat3 after injury***

a) *pSTAT3 expression in non-injured mouse cortex. Note the lack of expression in the absence of injury. (Scale bar 50µm).*

b) *Expression of pSTAT3 and Iba1 expression 24h after injury. Yellow arrows mark Iba1 cells that are also pSTAT3-positive cells. Green arrows mark Iba1 cells that are negative for pSTAT3. Note the reduction in Iba1/pSTAT3 positive cells in Sema4B knockout mice. (Scale bar 50µm).*

c) *Expression of pSTAT3 and S100β expression 24h after injury. Yellow arrows mark S100β cells which are also pSTAT3-positive cells. Green arrows mark pSTAT3-positive cells that are also negative for S100β cells. White lines point toward double-positive cells. Note that there is no significant change in S100 β/pSTAT3 cells in Sema4B knockout mice (Scale bar 50µm).*

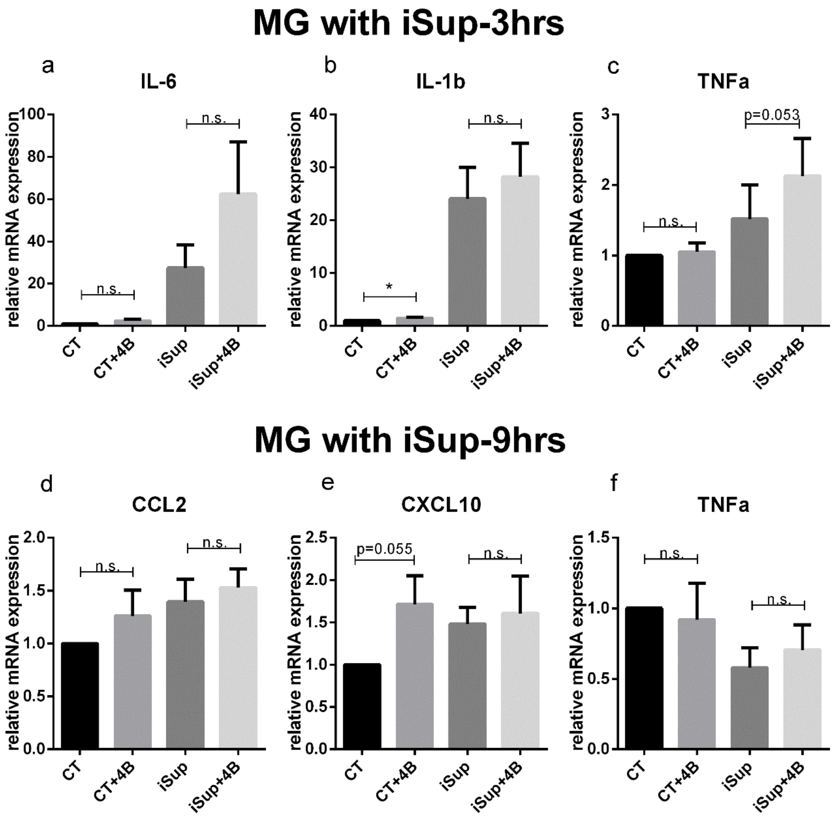


***Figure S7: Sema4B and iSup activation of cultured microglia***

*a-c) qPCR analysis of cultured microglia treated with Sema4B and/or iSup for 3h (n=6-9; Wilcoxon one-tailed test).*

*d-f) qPCR analysis of cultured microglia treated with Sema4B and/or iSup for 9h (n=7-10; Wilcoxon one-tailed test).*

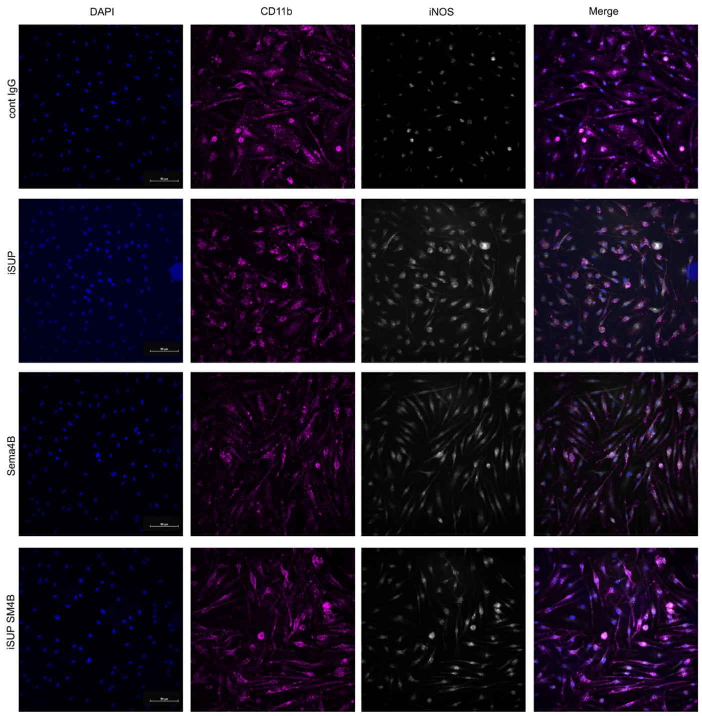


*Figure S8: iNOS expression in cultured microglia*

*Representative images of iNOS (gray) and CD11b (magenta) staining of microglial cultured cells expressing iNOS 24h after treatment (Scale bar 50μm).*

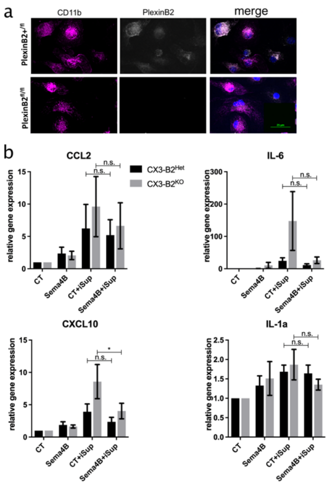


***Figure S9: activation of Cx3cr1creER:PlexinB2$^{fl/fl}$ microglia in culture***

Deleted: *8*

a) *Microglia cells were treated with OH-TAM. 10 days later the cells were statin with anti-PlexinB2 and CD11b (Scale bar 25µm).*

b) *) qPCR analysis of cultured microglia treated with Sema4B and/or iSup for 3h (n=6-9; Wilcoxon one-tailed test).*

**Supplementary methods:**

**Animal information**

All cortical injury experiments were performed on mice aged 7−12 weeks old. Genotype was determined by PCR analysis of genomic DNA isolated from tail clippings of 3-week-old mice. The presence of the wild-type Sema4B allele was established using primer 1 (5'-AGACATGGTGCTGGAGAGGT-3') with primer 2 (5'-TGTGTTTGGTTGGATCTGGA-3'). The mutant allele of Sema4B$^{-/-}$ was verified with primer 3 (5'-TGCACATGCTTTACGTGTG-3') and primer 4 (5'-TGCCGCGTGTCGTGTTGCAC-3'). For Cx3cr1creER:PlexinB2$^{fl/fl}$ mice: The presence of Cre was established using primer 5 (5'ATTTGCCTGCATTACCGGTC -3') with primer 6 (5'ATCAACGTTTTCTTTTCGG-3'). The PlexinB2 flox site was established using primer 7 (5'ACGTCATTCGTCTGGTCCTC-3') and primer 8 (5'GCTGCAAGAAGGAATTCACA-3').

**Stab wound injury model**

For the injury experiments, mice were anesthetized with a ketamine/xylazine solution (50 mg/kg ketamine/7.5 mg/kg xylazine in 0.9% NaCl solution). A sterile needle (1.2mm) was inserted vertically into the right cerebral hemisphere, reaching the skull surface at a depth of 5mm. The needle was inserted through the cranium 2mm caudal to the bregma and 1mm lateral to the midline. The skin incision was closed with biological glue.

**Tamoxifen treatment:**

Tamoxifen (Cayman Chemical Company, #13258) dissolved in 5% ethanol and corn oil at a concentration of 20mg/ml was injected intraperitoneally into mice for 4 days, 125µl/mouse, and stab injuries were performed 3 weeks after the end of treatment. 4-OH tamoxifen (1µM) dissolved in 100% ethanol was added to microglial cultures for 5 days before the experiment was performed.

**Fluoro-Jade C staining**

Frozen sections were fixed with 4% PFA, washed 3 times for 5 minutes each with PBS, and dried at 60°C for 30 min. Slides were immersed in 100% ethanol for 5 min, 70% ethanol for 2 min, and DDW for 2 min. They were then incubated with 0.06 % potassium permanganate for 15 min, followed by immersion in DDW for 2 min. They were incubated with 0.0001% FJC (Histo-Chem Inc., Jefferson, Ar, USA) in 0.1% acetic acid for 20 min, and again immersed in DDW 3 times for 1 min each. The slides were dried again at 60°C for 5 min and then incubated with Xylene for another 5 min. Finally, mounting medium DPX (Sigma #44581) was applied, and the slides were covered with coverslips. The staining was analyzed using fluorescent microscopy (Ex 435, Em 525).

**RNA extraction and qPCR:**

To preserve RNA integrity, a triazole-based RNA KIT-Direct-zol™ RNA MiniPrep (Zymo Research) kit was used and the acquired RNA was always placed on ice or at -80°C degrees. cDNA was prepared using qScript cDNA Synthesis Kit (Quantabio).

qPCR was performed with triplicates per sample, using Universal SYBR Green Supermix (BioRad, Hercules, CA). Differential expression was determined using the delta CT method. Each primer set was tested on human and mouse samples to make sure each primer is mouse specific. The primers used are as follows:

qPCR primers:

| gene | Fw seq | Rev seq |
|---|---|---|
| Snrpd3 | TTCGCCTTCTAACGTTTGTG | CATCTTGGCAGGACTCTTCC |
| Chmp2A | AAGGCCAGATGGATGCTGT | CCGCATCAACACAAACTTGC |
| CCL2 | AGGTCCCTGTCATGCTTCTG | TCATTGGGATCATCTTGCTG |
| CCL3 | CCCAGGTCTCTTTGGAGTCA | AGATTCCACGCCAATTCATC |
| IL-6 | GTTCTCTGGGAAATCGTGGA | GGTACTCCAGAAGACCAGAGGA |
| CCL5 | CCCACTTCTTCTCTGGGTTG | GTGCCCACGTCAAGGAGTAT |
| ATF3 | CCCCTGGAGATGTCAGTCAC | GCAGGCACTCTGTCTTCTCC |
| Kklhl6 | GCTGAGCTGTGTCACCAGAC | AGCCATGCCAGTAGAGGCTA |

| | | |
|---|---|---|
| Txnip | GGTCTCAGCAGTGCAAACAG | AGCTCGAAGCCGAACTTGTA |
| Cadm3 | CTGCCTTCCTTTACCCACAC | TTAGGCCAGAACCTGCTCAT |
| GPR88 | CTGGCCAACTCTTCACACCT | CTAAGGACTGGCCCCAAATC |
| CD200 | CTCCTGATTTCGGTGACGTT | CCTGGGGAATGTGATTGACT |
| IL-1α | AGAGAGATGGTCAATGGCAGA | TCAAGATGGCCAAAGTTCCT |
| CXCL10 | CCTATGGCCCTCATTCTCAC | AAGTGCTGCCGTCATTTTCT |
| IL-1β | ACCTTCCAGGATGAGGACATGA | CTAATGGGAACGTCACACACCA |
| TNFα | CCACATCTCCCTCCAGAAAA | GTGGGTGAGGAGCACGTAGT |
| PlexinB1 | CTGATACCGGTCCATGTGGAACGC | GGAAGCTGGGTCCTGAAGGCTG |
| PlexinB2 | CCCAAGCAGCATCCCTAGTA | GGAGCACAGCTACAGTGTGTCAC |
| Chil3 | AGACCTCAGTGGCTCCTTCA | TAGTACTGGCCCACCAGGAA |
| Psmb8 | CAGTCCTGAAGAGGCCTACG | CACTTTCACCCAACCGTCTT |

**Immunofluorescence analysis**

Brains were fixed in 4% PFA, followed by incubation in 30% sucrose for at least 24h and then the brains were frozen in OCT (Tissue Tek #4583) in liquid nitrogen. Coronal tissue sections (20µm thick) were fixed again in 4% PFA, washed 3 times for 5 minutes each, and then incubated for 1h at room temperature in a blocking solution consisting of 0.5% Triton X-100, 0.01% $NaN_3$, and either 5% goat serum, FBS, or BSA. This solution was used for the dilution of both primary and secondary antibodies. Sections were incubated in the primary antibody overnight (16h). Slides were then washed three times (5 min each) with PBS and visualized with Cy3- and Cy5-labeled secondary antibodies (Jackson Immuno Research Laboratories, Inc.) and covered with fluorescent mounting media (Dako). Sections were photographed using a spinning disc fluorescent microscope (Nikon) and analyses were done by ImageJ software.

**Antibody list:**

| antibodies | Species | Company | Cat # | Dilution |
|---|---|---|---|---|
| Iba1 | rabbit | Biocare Medical | CP290B | 1:500 |

| | | | | |
|---|---|---|---|---|
| Iba1 | goat | Novus Bio | NB100-1028 | 1:700 |
| Tmem119 | rabbit | abcam | ab209064 | 1:800 |
| CD45 | Rat | Invitrogen | 14-0451-82 | 1:250 |
| pSTAT3 | rabbit | Cell Signaling Technology | 9145 | 1:100 |
| CD11b | Rat | Developmental Studies Hybridoma Bank | M1/70.15.11.5.2 | 1:20 |
| Aldh1L1 | Rat | Developmental Studies Hybridoma Bank | N103/31 | ? |
| iNOS | Rabbit | Abcam | ab15323 | 1:250 |
| PlexinB2 | hamster | ebioscience | 14-5665-82 | 1:500 |
| F4/80 | Rat | abcam | ab6640 | 1:500 |
| S100β | mouse | abcam | ab11178 | 1:400 |

**List of antibodies used in the panning experiments:**

| **a**ntibody | **c**oncentrations | **Catalog number** | **c**ompany |
|---|---|---|---|
| Goat Anti-Mouse IgG + IgM (H+L) | 0.5µg/ml | (**1**15-005-044) | Jackson Immunoresearch |
| Anti-CD45 Monoclonal Antibody | 0.5µg/ml | (30-F11) | eBioscience™ |
| anti-CD90/Thy1 MAb | 0.5µg/ml | MAB7335 | R&D systems |

| | | | |
|---|---|---|---|
| Anti-O4 MAb | 0.5µg/ml | (Clone O4), MAB1326 | R&D systems |
| Anti-ACSA-2 | 0.1 µg/ml | 130-099-138 | Miltenyibiotec |

**Sema4B and Fc Fusion Proteins preparation:**

The Fc fusion protein of the ectodomain of mouse Sema4B (amino acid 1-700) was cloned into PsxFc2 vector upstream of the hinge region of human IgG1-Fc. Fc protein was used as a control. 2.5 million HEK293T cells were transfected with 12µg DNA in 350µm of DMEM only and 25µl of Linear-Polyethylenimine reagent (Polysciences, Warrington, PA, USA) per 10cm$^2$ plate. The day after transfection the medium was changed to DMEM only. The concentration of the recombinant proteins was quantified by Western blot using anti-hIgG secondary Ab. Known quantities of IgG from human serum were used to estimate the amount of Fc fusion proteins. After blocking for 1h, and one wash with TBST for 5 min, the membrane was incubated with peroxidase-conjugated anti-Human IgG ab (Jackson ImmunoResearch #109-035-003 ,1:1000 dilution) for 1h, and then developed. The quantification of concentrations was performed using ImageJ software.

**Astrocyte and microglia cultures:**

Cerebral hemispheres were aseptically removed from newborn (1-3 day-old) pups, the meninges were removed, and the cortices were incubated in 0.25% Trypsin B for 10 min at 37°C. Tissues were then mechanically dissociated by trituration in Dulbecco's modified Eagle's medium (DMEM) high glucose containing 10% fetal bovine serum (FBS), 2% AmphoB, 1% each of P/S, L-Glu, Hepes, and sodium pyruvate. They were then centrifuged at 300g for 5 minutes, the supernatant was removed, and they were resuspended in a medium. For mixed cell cultures, we plated the cortices from 3 brains in a 10cm$^2$ plate, or 1 brain in a 30mm$^2$ well. The cultures were thoroughly washed with PBS after 2-3 days and the medium was replaced with DMEM complete (DMEM high glucose, 10% FBS, 1% each of P/S, L-Glu, Hepes, & AmphoB) and GM-CSF (5ng/ml). Astrocyte cultures were filtered through a 40um filter before centrifugation and the cortices from one brain were plated in a T75 flask with EGF (20ng/ml). The astrocytes were thoroughly washed with PBS after 2-3 days, and the medium was replaced with DMEM complete with another round of EGF. Upon confluence, the medium was replaced with D-Valine medium

(0.96% D-Val powder, 10% Dialyzed FBS, 2% Hepes, & 1% each of P/S & AmphoB in DDW, pH 7.4) and 1mM AraC. After 2 days, the medium was replaced with DMEM complete, and the astrocytes were put in shaking overnight at 37°C. The next day, the astrocytes were passaged 1:3 using TrypLE. AraC was added again upon confluence, and the cells were cultured for the experiment 3 days later. Cells were cultured on poly-D-lysine treated plates or flasks (50µg/ml at 37°C for 0.5h, with 3 washes of DDW for 5 minutes each, and 0.5h of UV radiation). Microglia were purified from mixed cell cultures by adding 2.5mM of Lidocaine (Sigma L5647), incubating for 15 minutes at room temperature, then centrifuging the collected medium for 5 minutes at 500rcf with 50mM of EDTA (pH 8.0). The supernatant was removed, and cells were resuspended in a lower volume of DMEM complete.

All medium was filtered through 0.22um filters before use.

Reagents: PDL (Sigma, #P1024). Biological industries: Trypsin B 0.25% (#03-046-1B), L-Glu (#03-020-1B), P/S (#03-031-1B), sodium pyruvate 100mM (#03-042-1B), DMEM (#01-055-1A). D-Val (#M38861-20, US Biological), PBS without Mg++ and Ca++ (BP655/500D, Hylabs), HEPES 1M, pH 7.5 (Sigma), AmphoB 2.5mg/ml (#11636, Cayman Chemicals), FBS (#SV30160.03, Lot #RE00000002, Hyclone), TrypLE (#12604-021, Gibco), GM-CSF (Reprokine, #RCP01587), EGF (Reprokine, RKP01133).

**COS cell collapse assay:**

COS-7 cells were maintained in Dulbecco's's modified Eagle's medium supplemented with 10% fetal bovine serum. Cells were transfected with 1µg DNA. In all experiments, cells were transfected with expression vectors for pEGFP (200ng) and PlexinB2 (800ng) or carrier DNA (pBS plasmid). Linear PEI (Polysciences) was used as a transfection reagent in all experiments. Thirty-six fields of cells were photographed using a GFP channel 48 hours after transfection, to monitor only the transfected cells. The same fields were photographed once more 40 min after the addition of Sema4B (50ng/ml). To scan the same fields of cells we used a scanning stage apparatus. Cell images were captured using a fluorescence microscope (Olympus, Hamburg, Germany) equipped with a cooled CCD camera (Roper Scientific, Duluth, GA, USA) and a 20x objective. The cells were counted and classified according to their morphology by an observer who was blind to the identities of the treatments.
